# Mutational analysis of the nitrogenase carbon monoxide protective protein CowN reveals that a conserved C-terminal glutamic acid residue is necessary for its activity

**DOI:** 10.1101/2023.08.25.553292

**Authors:** Dustin L. Willard, Joshuah J. Arellano, Mitch Underdahl, Terrence M. Lee, Avinash S. Ramaswamy, Gabriella Fumes, Agatha Kliman, Emily Y. Wong, Cedric P. Owens

## Abstract

Nitrogenase is the only enzyme that catalyzes the reduction of nitrogen gas into ammonia. Nitrogenase is tightly inhibited by the environmental gas carbon monoxide (CO). Many nitrogen fixing bacteria protect nitrogenase from CO inhibition using the protective protein CowN. This work demonstrates that a conserved glutamic acid residue near CowN’s C-terminus is necessary for its function. Mutation of the glutamic acid residue abolishes both CowN’s protection against CO inhibition and CowN’s ability to bind to nitrogenase. In contrast, a conserved C-terminal cysteine residue is not important for CO protection. Overall, this work uncovers structural features in CowN that are required for its function and provides new insights into its nitrogenase binding and CO protection mechanism.

## MAIN TEXT

Nitrogenase is the only enzyme capable of reducing dinitrogen gas (N_2_) into ammonia (NH_3_). It consists of two components proteins, the molybdenum iron protein (MoFeP) and the iron protein (FeP). MoFeP contains two metal clusters, the FeMoco, a [7Fe:9S:1Mo:1C] cluster that is the site of catalysis, and the [8Fe:7S] P-cluste.^*1, 2*^ FeP is a reductase that contains a [4Fe:4S] cluster. During nitrogen reduction, FeP sequentially delivers electron to MoFeP in an ATP-dependent process (Equation 1).

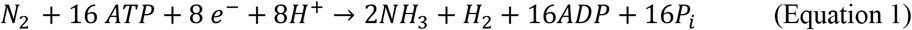

Molybdenum nitrogenase, the most common isoform of the enzyme, is inactivated by the environmental gas carbon monoxide (CO), which acts as a mixed inhibitor.^*3, 4*^ CO is produced anthropogenically during the combustion of fossil fuels, through natural geochemical and biological processes in the soil,^*5-7*^ and by plants, which use it is a signaling gas.^*8, 9*^ While CO potently inhibits nitrogenase *in vitro*, diazotrophs tolerate the gas quite well. The inhibition constants, *K*_is_ and K_*ii*_, are in the range of 1-2 × 10^−4^ atm and 1-2 × 10^−3^ atm, respectively, *in vitro* ^*3, 10, 11*^. In contrast, when nitrogenase activity is measured *in vivo* in diazotrophs the constants are an order of magnitude higher.^*3*^

In many α-proteobacteria, nitrogenase is protected by a small protein called CowN. *Rhodobacter capsulatus*^*12*^ and *Rhodospirillum rubrum*^*13*^ are able to grow diaoztrophically in 1% and 0.15% (0.01 atm and 0.0015 atm) CO, respectively, without apparent growth inhibition. However, when *cowN* in knocked out in either bacterium, diazotophic growth is nearly nonexistant, suggesting that *cowN* is vital in maintaining nitrogenase activity under CO. Our group recently described the mechanism of CowN protection in the diazotroph *Gluconacetobacter diazotrophicus* (*Gd*). CowN binds to MoFeP and increases the inhibition constant for CO-binding about 10-fold.^*10*^ There was no evidence that CowN alters the structure of FeMoco, nor does CowN have any activity of its own, leading to the hypothesis that CowN prevents CO from reaching FeMoco. CowN specifically prevents CO binding, as it does not prevent the alternative substrate acetylene (C_2_H_2_) from reaching the active site. While we were able to determine that CowN and MoFeP interact, the binding surface on either protein is unknown. This goal of this work is to identify structural features in CowN that are important for its function and for MoFeP binding. Mutagenesis and crosslinking data suggest that a glutamate residue near CowN’s C-terminus is necessary for CowN activity, however that a conserved cysteine residue is not essential for CO protection.

Our group previously examined nitrogenase protection by CowN using the common model substrate C_2_H_2_ .^*10*^ Since the mechanism of C_2_H_2_ and N_2_ reduction are different,^*14-16*^ we wanted to determine if CowN protects nitrogenase under N_2_ turnover. As shown in **Figure 1**, CowN prevents CO inhibition when N_2_ is the substrate. The amount of protection for N_2_ and C_2_H_2_ reduction is nearly identical, suggesting that CowN’s protective mechanism is likely the same for both gases. The result is consistent with our proposal that CowN blocks CO access to FeMoco and thus protects nitrogenase turnover for all substrates. Furthermore, these results directly explain the observations by Masepohl *et al*.,^*12*^ and Kerby *et al*.^*13*^ who demonstrated that CowN sustained diazotrophic growth in presence of CO.

**Figure 1.**
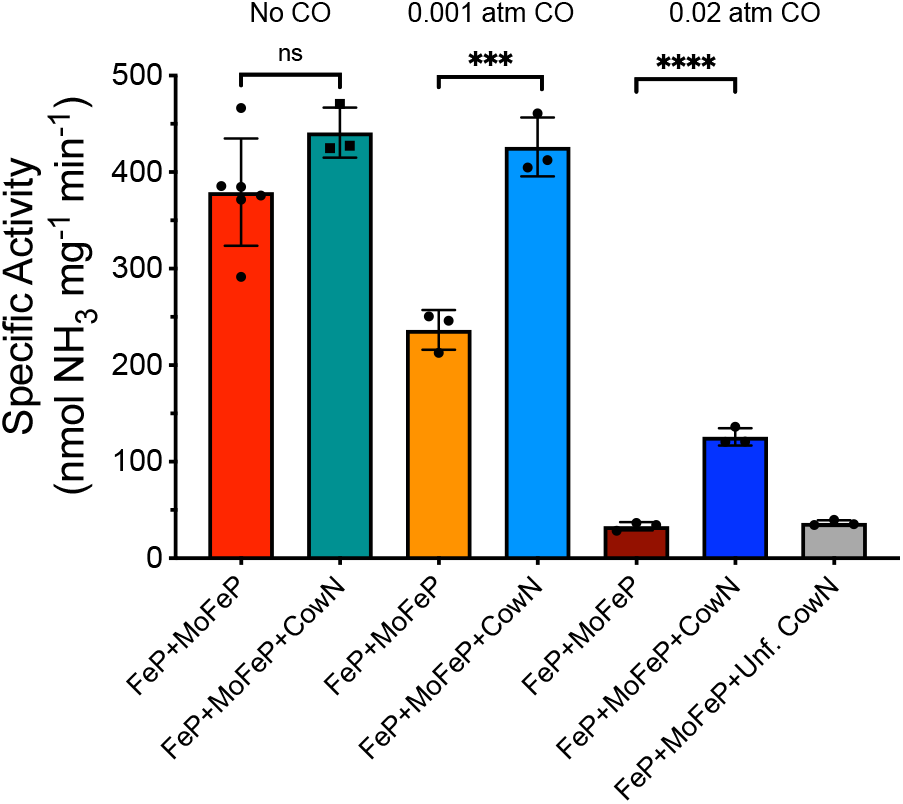
N_2_ reduction by nitrogenase under CO, with and without CowN. The stars represent results from an unpaired t-test comparing means, with *** meaning p < 0.01 and **** meaning p < 0.001.

CowN is predicted to be a compact, 13 kDa 4-helical bundle. CowN’s primary sequence is not particularly conserved, however, it commonly features a negatively charged C-terminal end that contains several Glu and Asp residues (**Figure 2A and 2B**). In *Gd*-CowN, the protein is polar with a negatively charged C-terminus and a more positively charged N-terminus (**Figure 2C**). A second conserved feature in CowN is the presence of a C-terminal sulfur containing residue (**Figure 2A and 2B**). CowN features a sulfur containing amino acid as either the penultimate or final residue. The residue is usually a Cys. In some cases, there are two thiol containing residues at the C-terminus, as is the case in *Gd*-CowN which ends in Cys90-Met91.

**Figure 2.**
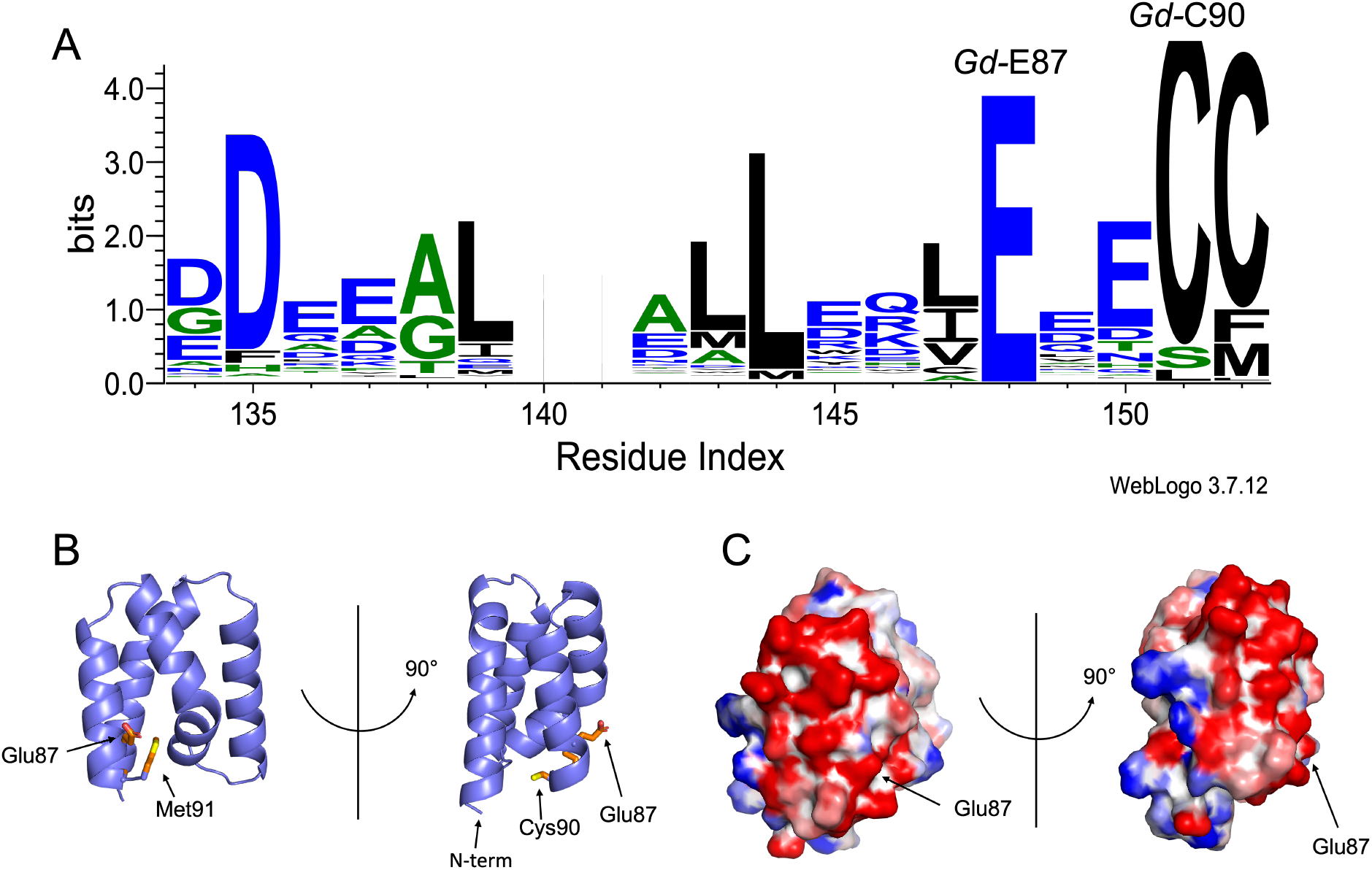
(A) Weblogos based on a Clusalω sequence alignment of the C-terminal end of all CowN sequences in Uniprot. The size of the amino acid is proportional to its conservation. The only fully conserved residue is Glu87 (*G. diazotrophicus* numbering). Furthermore, CowN nearly always (377 out of 378 sequences) features a Cys or a Met at either the ultimate or penultimate position. (B) Predicted Alphafold 2^*17*^ structure of *Gd*-CowN highlighting residues Glu87, Cys90, and Met91. (C) Calculated electrostatic map of *Gd*-CowN. The electrostatic map was generated using the Bluues server^*18*^ using default parameters. Only the 4-helical core of CowN (residues 13-91) is displayed as the unstructured N-terminal tail was deleted. Negative charges are colored red and positive charges are blue. The righthand view showcases the polarity of the protein. The views in (B) and (C) are identical.

The conservation of acidic and sulfur-containing residues at CowN’s C-terminus suggests functional relevance. To examine the importance of the negative C-terminus, we targeted Glu87, the only fully conserved residue in CowN, for mutagenesis, generating the E87A, E87D, and E87Q mutants. Furthermore, we investigated the role of the C-terminal Cys and Met residues by making a C90A, C90S, and M91A mutant as well as a C-terminally truncated version of CowN that lacks the final two residues (referred to as C-trunc CowN).

Mutant expression and purification were identical to WT CowN. All three E87 mutants and both C90 mutants expressed to similar levels as wild type, yielding approximately 3-4 mg protein per liter of growth media. Circular dichroism (CD) analysis suggests that the Glu87 and Cys90 mutants are folded α-helical proteins, as expected, (**Figure 2A**) and are identical to WT CowN. In contrast, M91A and C-trunc CowN expressed at lower levels (0.5-1 mg per liter of growth media). Most of the protein was present as a misfolded aggregate, however, a small amount of monomeric protein could be separated from the aggregate by gel filtration chromatography. CD spectra of monomeric M91A and C-trunc CowN indicates that the protein is less folded than wild-type CowN (**Figure 2B**). This suggests that residue Met91 is important for CowN stability. However, the reason for this is unclear. Met91 is not conserved. Structural modeling suggests that M91A CowN should be nearly identical to WT CowN (**Figure S1**). CowN has no predicted posttranslational modifications or ligand binding sites that would depend on Cys90 or Met91, and we did not detect modifications to either residue by mass spectrometry. Met91 is predicted to point towards the hydrophobic center of CowN (**Figure S1**). We therefore hypothesize that it may play a role in stabilizing hydrophobic packing between CowN’s α-helices.

To test the role of the conserved residue Glu87, we carried out turnover experiments with acetylene. These experiments were conducted with a CO concentration of 0.02 atm, which abolishes approximately 90% of nitrogenase activity in absence of CowN. Consistent with previous reports,^*10*^ WT CowN leads to a partial rescue of activity to 30% of the uninhibited level (**Figure 2C**). E87A, E87D, and E87Q CowN do not protect nitrogenase from CO. Their levels of protection are within error of samples that do not have CowN and of an unfolded CowN negative control. Additionally, we investigated if E87A is able to recover nitrogenase activity if it is present in higher concentrations (**Figure S2**). However, recovery increased only slightly when the E87A CowN concentration was increased to 4 μΜ (the highest amount that could be tested in our experimental setup), suggesting that the mutation is deleterious enough that it cannot be overcome by increasing the CowN concentration.

Both M91A and C-trunc CowN do not protect nitrogenase from CO inhibition (**Figure 3D**). This result was expected, given the fact that both mutants are partially unfolded. In contrast, both C90S and C90A CowN protect nitrogenase to the same degree as WT CowN (**Figure 3D**). Taken together, these data suggest that CowN function depends on residue Glu87 and that even mild substitutions to Gln and and Asp are not tolerated. At the same time, the data suggests that the conserved C-terminal Cys is not required for CO protection.

**Figure 3.**
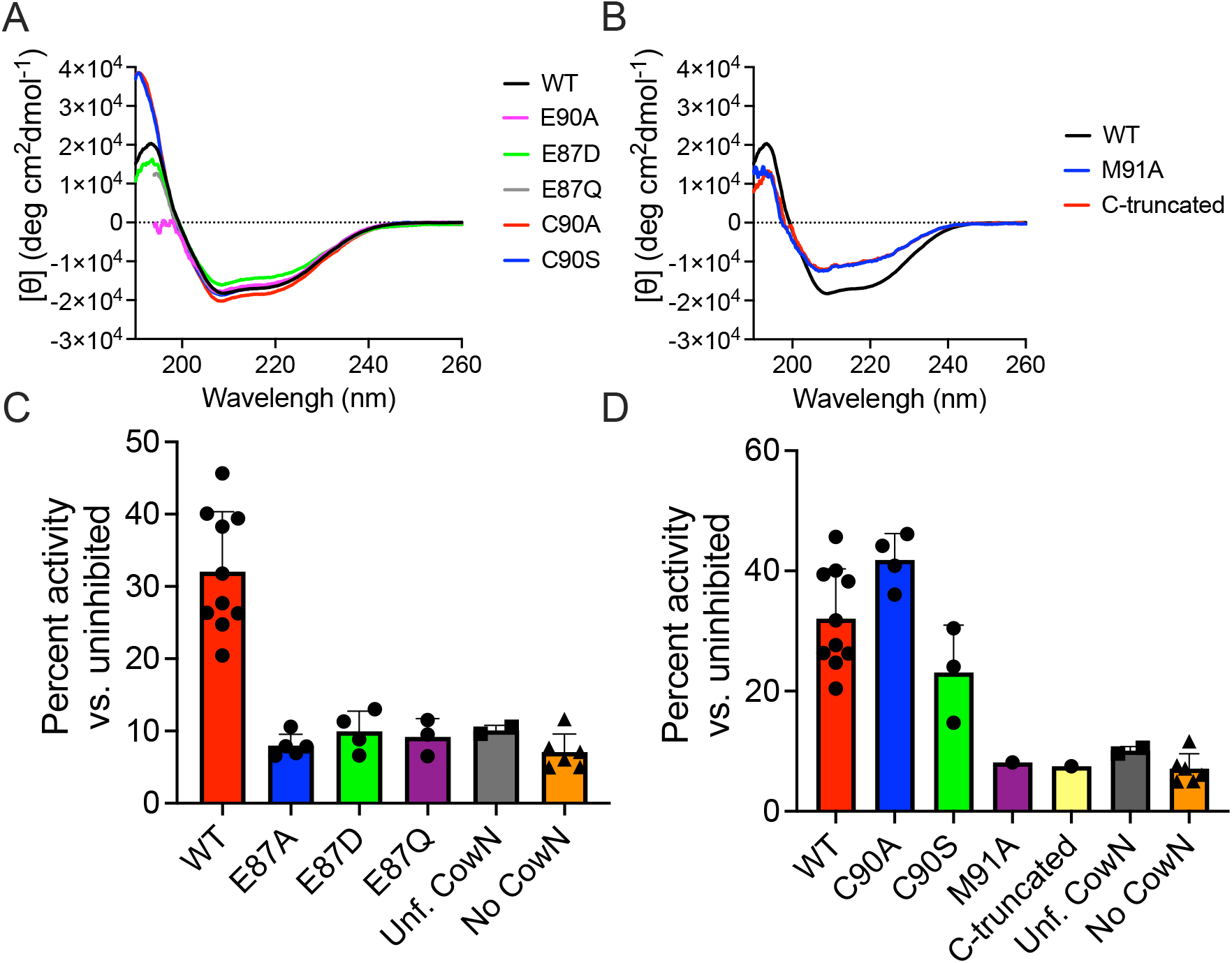
(A) CD spectra of the folded CowN mutants and (B) of the partially unfolded M91A and C-truncated variants. (C) CO protection of the Glu87 mutants and (B) Cys90 and Met91 mutants. Each point represents the average of an independent assay consisting of triplicate measurements. The difference in mean, as determined by an unpaired t-test, is statistically significant to a p-value of <0.01 between WT and E87A, E87D, E87Q, M91A, and C-truncated CowN. The differences in mean between WT CowN and C90A and C90S CowN is not significant. Unf. CowN stands for misfolded CowN oligomer. Percent recovery was determined by calculating the amount of activity relative to triplicate nitrogenase control samples that did not receive CO.

**Figure 4.**
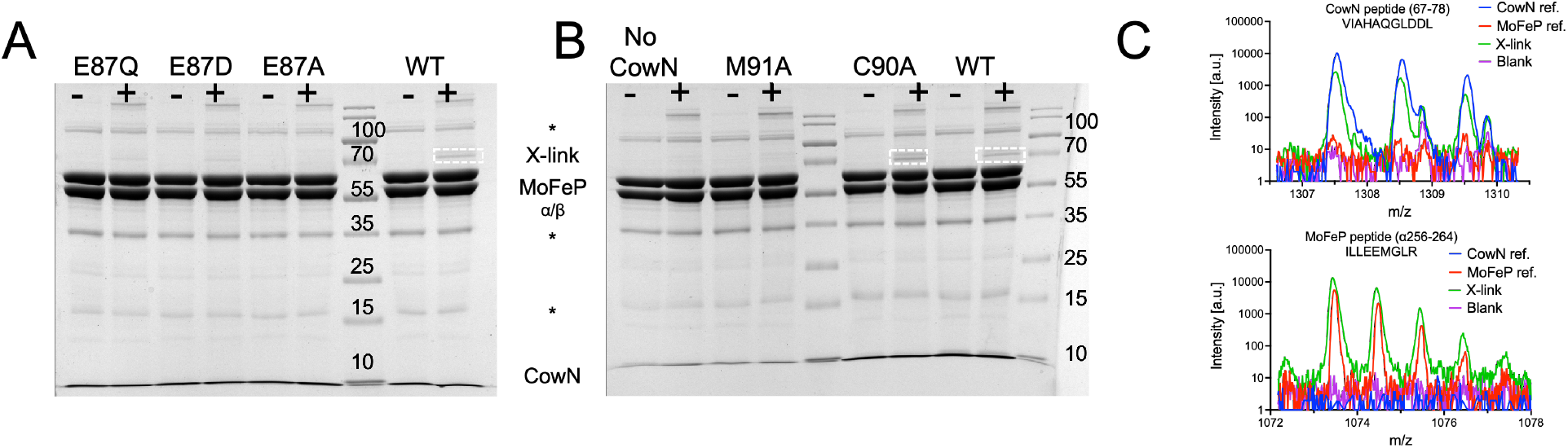
(A) SDS-PAGE of diazirine crosslinking between Glu87 CowN mutants with MoFeP and (B) C90A and M91A CowN mutants with MoFeP. The 70 kDa crosslinking band is highlighted with a dashed white box for clarity. (C) MALDI-TOF MS peaks after in gel trypsin digest illustrating the presence of characteristic CowN and MoFeP peptides in the excised crosslinking bands. MoFeP ref. and CowN ref. represent MoFeP and CowN-only controls for comparison. A “+” sign on the gels denotes samples that were illuminated. Asterisks indicate impurities in the MoFeP sample. The gels are representative examples of triplicate experiments.

We hypothesize that the loss of CO protection in the Glu87 mutants is caused by a decreased affinity for MoFeP, as Glu87 may form a salt bridge with a positively charged MoFeP residue. To test this hypothesis, we carried out crosslinking experiments with diazirine labeled MoFeP. In this experiment, MoFeP is first labeled at Lys residues with a photoactivatable diazirine crosslinker using NHS-ester chemistry. The diazirine crosslinker then binds covalently to interacting proteins under illumination. We had previously used this method to determine that MoFeP interacts with WT CowN.^*10*^ As shown in **Figure 3A and 3B**, WT CowN and MoFeP form a diazirine-mediated crosslink band at 70 kDa. A representative MoFeP and CowN peptide peak from an excised crosslink band that was tryptically digested and analyzed by mass spectrometry is shown **Figure 3C**. Furthermore, a list of peptides belonging to WT CowN and MoFeP from the excised crosslink band and a sequence coverage map are shown in **Tables S1-S3**. The crosslinking band intensity for E87D and E87Q CowN is weaker than for the wild type, and E87A CowN does not crosslink at all (**Figure 3A**). These data support the hypothesis that Glu87 is important in mediating protein-protein interactions with MoFeP. We also examined crosslinking of C90A and M91A CowN with diazirine-labeled MoFeP (**Figure 3B**). M91A did not crosslink with MoFeP, suggesting that its partially unfolded state prevents it from interacting with MoFeP. In contrast, C90A CowN bound to MoFeP, adding further evidence that mutations at residue Cys90 do not impair CowN function. Taken together, the crosslinking and turnover data are consistent with each other and suggest that residue Glu87 is important for CowN function as it is required for protein-protein interaction with MoFeP.

Glu87 likely enables CowN to bind to MoFeP through ionic interactions with positively charged residues on MoFeP. Such electrostatic interactions may occur at the opening of a putative CO channel between the α and β subunits, which is lined by four Lys residues.^*10, 14*^ This potential binding site is compatible with diazirine crosslinking results since the diazirine crosslinker is bound to MoFeP at a Lys residue. While the hypothesis that CowN blocks the mouth of a putative CO channel makes mechanistic sense, further work is needed to confirm this site. We carried out additional mass spectrometry experiments with different mass detectors and using methods for finding large peptides, however, we were unable to find peaks corresponding to crosslinked peptide pairs. The most likely explanation for the lack of detection is the low abundance of crosslinked peptide fragments and their large size. A possible solution to overcome this problem is to utilize a cleavable heterobifunctional diazirine crosslinker to detect binding sites using MS^3^ methods.^*19, 20*^ However, such crosslinkers are not yet commercially available.

Overall, the data presented herein suggests that CowN’s negatively charged C-terminal is important for CowN binding to MoFeP and protection against CO inhibition. At the same time, this work raises the question about the role of the C-terminal Cys. We previously speculated, that CowN may have additional roles in nitrogen fixation that are unrelated to CO protection since CowN is upregulated under all nitrogen fixation conditions, not just when CO is present.^*10*^ Independently, other researchers have implicated CowN in nitrogenase cold protection.^*21, 22*^ Our group intends on investigating CowN further to uncover the full gamut of its roles in bacterial nitrogen fixation and to unambiguously determine where it interacts with MoFeP.

## ASSOCIATED CONTENT

### Supporting Information

The supporting information contains detailed experimental methods, a structural model of M91A CowN, data on E87A CowN CO protection at higher protein concentration, and tables listing the peptides found by mass spectrometry.

### Accession numbers

*G. diazotrophicus* CowN Uniprot ID: A9H4F6

### Author Contributions

Experimental design, execution and analysis: Dustin L. Willard, Joshuah J. Arellano, Mitch Underdahl, Terrence M. Lee, Avinash S. Ramaswamy, Emily Y. Wong, Gabriella Fumes, Agatha Kliman, Cedric P. Owens; Funding acquisition, Cedric P. Owens; Writing and editing: Cedric P. Owens; All authors have given approval to the final version of the manuscript.

### Funding Sources

This work was supported by National Science Foundation, Division of Chemistry grant 1905399 and a Research Corporation for Science Advancement Cottrell Scholar Award to C.P.O.

## ACKNOWLEDGMENT

The authors would like to acknowledge Mr. Benjamin Katz and the University of California, Irvine mass spectrometry facility for help with MALDI-TOF mass spectrometry experiments. We also thank Drs. Bisoffi and Fudge for use of instrumentation.

## ABBREVIATIONS

BS3: (Sulfo-DSS) is bis(sulfosuccinimidyl)suberate
EDC: (1-ethyl-3-(3-dimethylaminopropyl)carbodiimide hydrochloride)
ESI: Electrospray ionization
FeMoco: iron-molybdenum cofactor
FeP: iron-protein
MALDI-TOF: Matrix assisted laser desorption ionization-time of flight
MoFeP: molybdenum-iron protein
SIAB: (succinimidyl (4-iodoacetyl)aminobenzoate)

