## Supporting information for "Mutational analysis of the nitrogenase carbon monoxide protective protein CowN reveals that a conserved C-terminal glutamic acid residue is necessary for its activity"

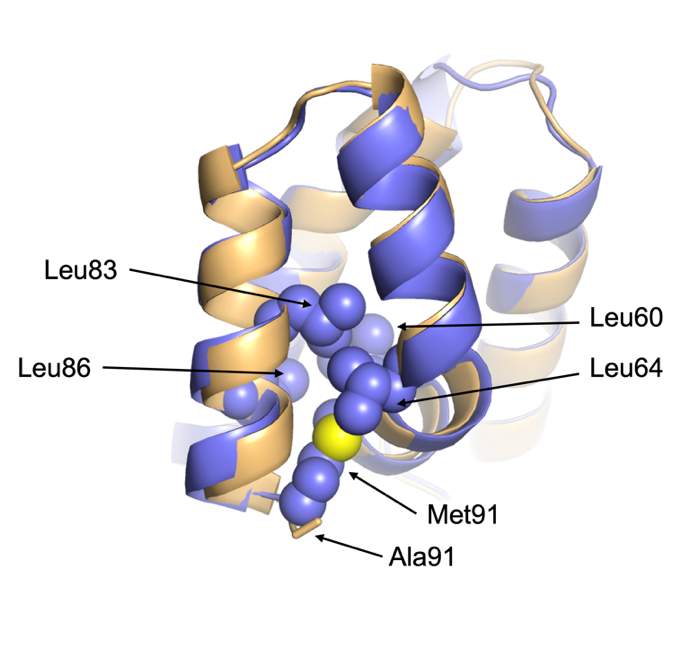
­­

**Figure S1.** Structural model of M91A CowN (orange) aligned to a model of WT CowN (blue), showing that M91A CowN is predicted to be identical to WT CowN. Both models were generated using Robetta.*^4^* In WT CowN, Met91 may be involved in forming a hydrophobic core with three leucine residues, Leu60, Leu64, Leu83, and Leu86. Ala91 would not be able to take part in such hydrophobic packing.

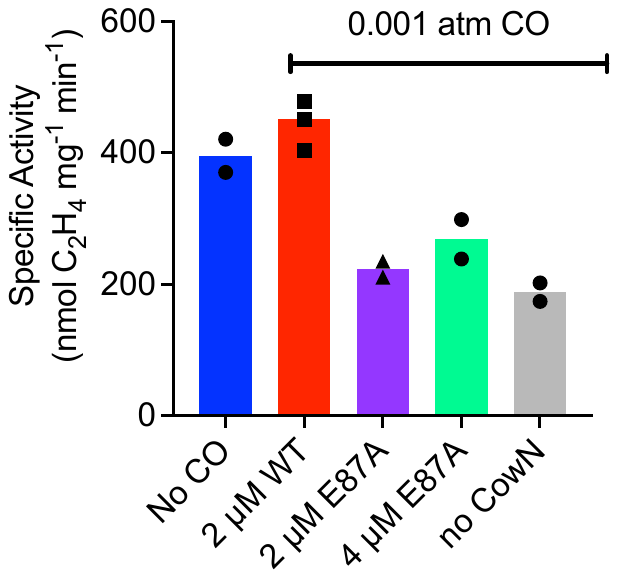

**Figure S2.** C_2_H_2_ reduction by nitrogenase under 0.001 CO with E87A CowN. The protein concentrations for MoFeP and FeP were 0.2 µM, 2 µM, respectively. E87A CowN was at either 2 µM or 4 µM, as indicated. The CO concentration was 0.001 atm. At this concentration, nitrogenase is approximately 50% inhibited in absence of CowN. We chose to use 0.001 atm CO in this experiment compared to 0.02 atm, which is what was used in the main text, since we expected to observe a greater amount of recovery when lower CO levels are used. For comparison, WT CowN confers full recovery under these conditions. However, as indicated, E87A CowN does not enhance nitrogenase activity substantially, even with lower CO levels and at higher protein concentration.

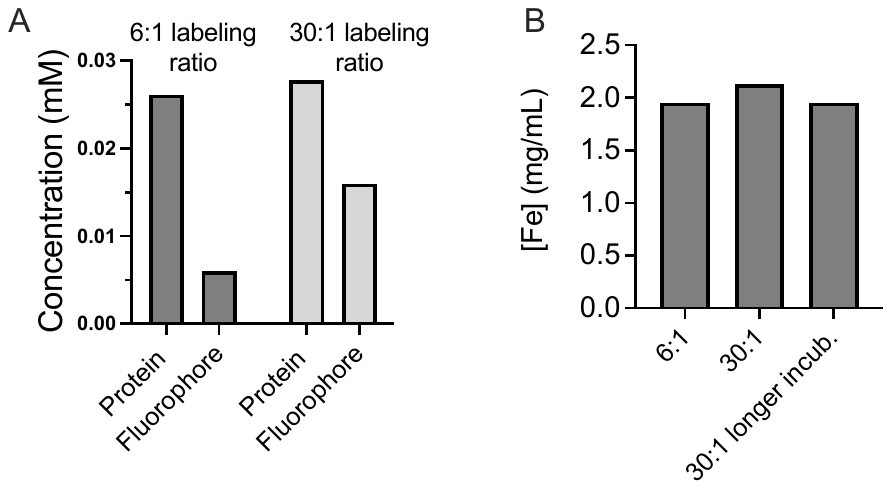

**Figure S3.** (A) Labeling MoFeP with 5-iodoatemido fluorescein, indicating that the degree of labeling scales with the amount of excess label that was added to MoFeP. At the 30:1 ratio, which is equivalent to the amount of excess SIAB that was used in subsequent experiments, there is one fluorescein label per MoFeP tetramer. Protein concentration was determined using the Bradford assay and the fluorophore concentration calculated using ε_494 nm_ = 80 mM^-1^cm^-1^ (B) Metal content analysis of labeled MoFeP samples demonstrates that adding more label does not cause to Fe loss. The 6:1 and 30:1 samples were incubated for 30 min, whereas the “30:1 longer incub.” sample was incubated for 60 min, suggesting that longer incubation does not damage the metal clusters either. The data in (A) and (B) represent the averages of duplicate independent measurements. Iron contents was determined using the 2,2 bypridine iron chelation assay, as described in Medina et al.*^1^*

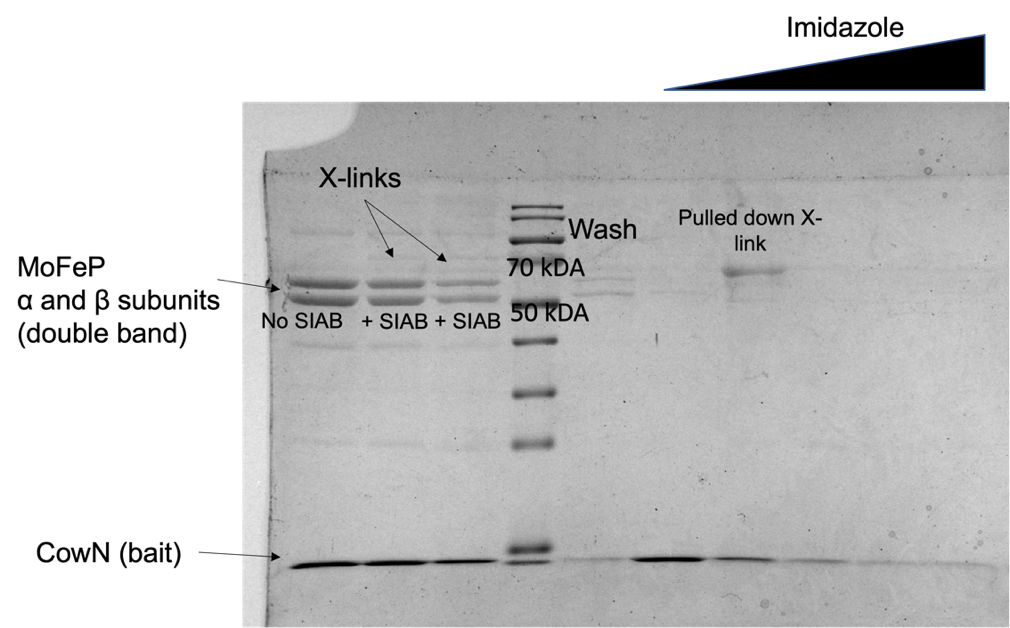

**Figure S4.** Example of a pulled down CowN-MoFeP complex. The protein concentrations in the crosslinking experiments were 1.5 µM MoFeP and 15 µM CowN. The reaction volume was 5 mL. His-tagged CowN served as the bait to pull down crosslinked CowN-MoFeP complex. Proteins were pulled out of solution using Ni-NTA beads. The pulldown occurred under denaturing conditions with 6M Urea since MoFeP otherwise sticks to the Ni-NTA resin. The pulled down crosslink band was trypsinized and analyzed by LC-MS/MS. Although there was excellent sequence coverage for MoFeP and CowN, no crosslinked peptides were found.

**Table S1. Summary of crosslinking results listing the number of unique peptides identified in different experiments**

| **Experiment**  **No. of**  **matched peptides** | **SIAB X-linking with WT-CowN** | **SIAB X-linking and pulldown with WT-CowN** | **SIAB X-linking with C90A CowN** |
| --- | --- | --- | --- |
| CowN | 6 | 12 | 6 |
| α-subunit | 36 | 25 | 36 |
| β-subunit | 29 | 26 | 34 |

**Table S2. Coverage map and list of CowN peptides identified in SIAB X-linking experiments between WT CowN and MoFeP**

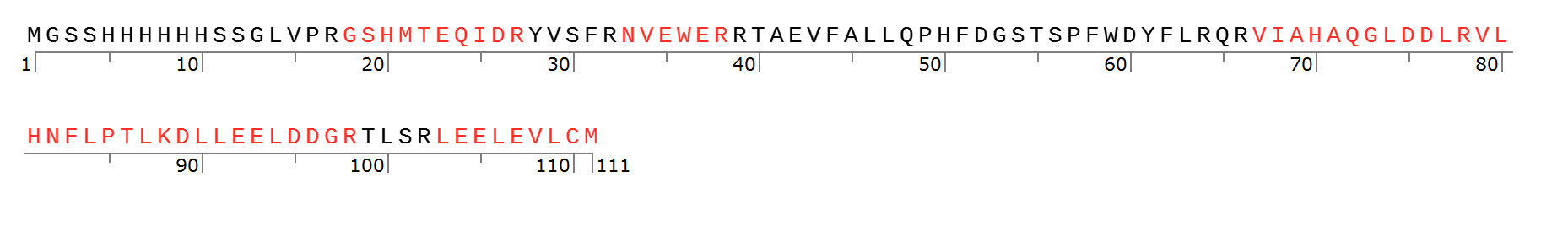

Six peptides identified for a total coverage of 51.4%

DLLEELDDGR

GSHMTEQIDR

LEELEVLCM

NVEWER

VIAHAQGLDDLR

VLHNFLPTLK

**Table S3. Coverage map and list of MoFeP α-subunit peptides identified in SIAB X-linking experiments between WT CowN and MoFeP**

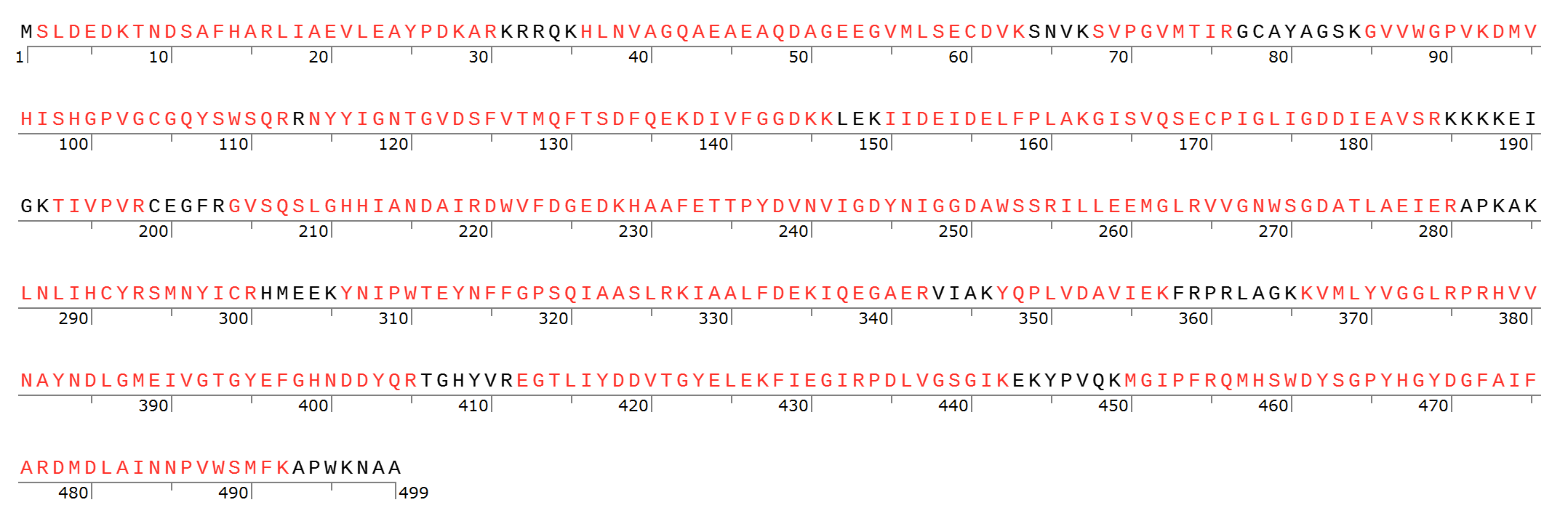

36 peptides identified for a total coverage of 84.6%

| DIVFGGDK | KIAALFDEK |
| --- | --- |
| DIVFGGDKK | KVMLYVGGLR |
| DMDLAINNPVWSMFK | KVMLYVGGLRPR |
| DMVHISHGPVGCGQYSWSQR | LIAEVLEAYPDK |
| DWVFDGEDK | LIAEVLEAYPDKAR |
| EGTLIYDDVTGYELEK | LNLIHCYR |
| FIEGIRPDLVGSGIK | MGIPFR |
| GISVQSECPIGLIGDDIEAVSR | NYYIGNTGVDSFVTMQFTSDFQEK |
| GVSQSLGHHIANDAIR | QMHSWDYSGPYHGYDGFAIFAR |
| GVVWGPVK | SLDEDKTNDSAFHAR |
| GVVWGPVKDMVHISHGPVGCGQYSWSQR | SMNYICR |
| HAAFETTPYDVNVIGDYNIGGDAWSSR | SVPGVMTIR |
| HLNVAGQAEAEAQDAGEEGVMLSECDVK | TIVPVR |
| HVVNAYNDLGMEIVGTGYEFGHNDDYQR | VMLYVGGLR |
| IAALFDEK | VMLYVGGLRPR |
| IAALFDEKIQEGAER | VVGNWSGDATLAEIER |
| IIDEIDELFPLAK | YNIPWTEYNFFGPSQIAASLR |
| ILLEEMGLR | YQPLVDAVIEK |

**Table S4. Coverage map and list of MoFeP β-subunit peptides identified in SIAB X-linking experiments between WT CowN and MoFeP**

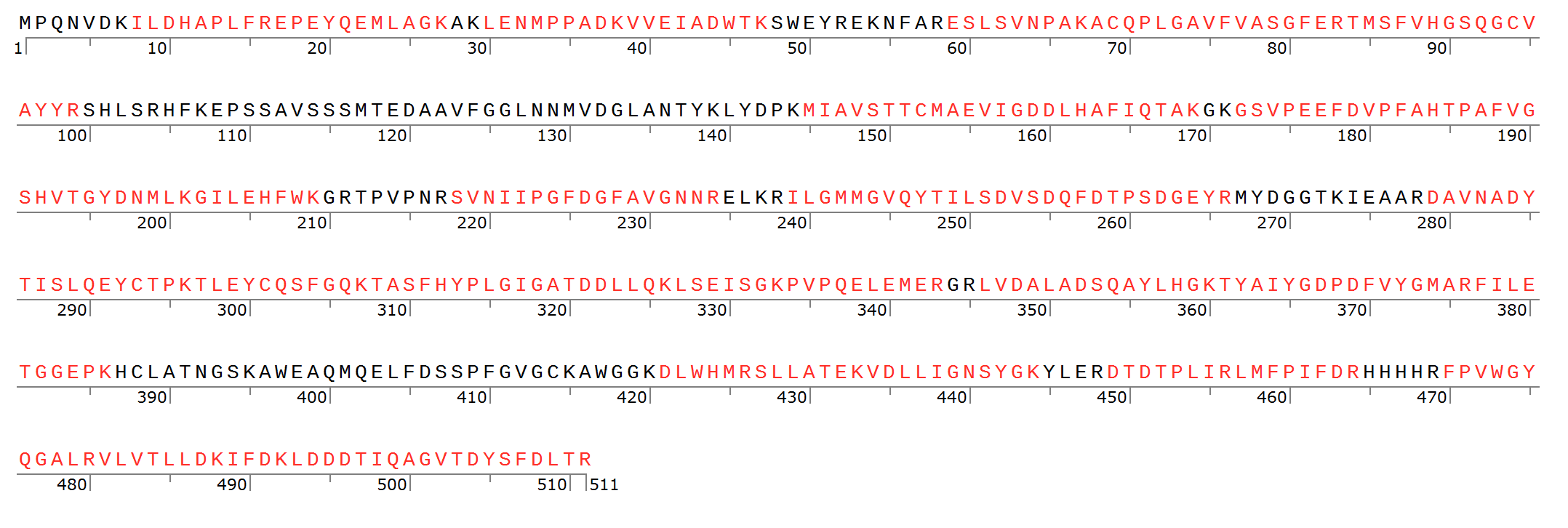

26 peptides identified for a total coverage of 77.3%

| ACQPLGAVFVASGFER | LENMPPADK |
| --- | --- |
| AWEAQMQELFDSSPFGVGCK | LMFPIFDR |
| DAVNADYTISLQEYCTPK | LSEISGKPVPQELEMER |
| DLWHMR | LVDALADSQAYLHGK |
| DTDTPLIR | MIAVSTTCMAEVIGDDLHAFIQTAK |
| EPEYQEMLAGK | SLLATEK |
| ESLSVNPAK | SVNIIPGFDGFAVGNNR |
| FILETGGEPK | TASFHYPLGIGATDDLLQK |
| FPVWGYQGALR | TLEYCQSFGQK |
| GILEHFWK | TMSFVHGSQGCVAYYR |
| GSVPEEFDVPFAHTPAFVGSHVTGYDNMLK | TYAIYGDPDFVYGMAR |
| IFDKLDDDTIQAGVTDYSFDLTR | VDLLIGNSYGK |
| ILDHAPLFR | VLVTLLDK |
| ILGMMGVQYTILSDVSDQFDTPSDGEYR | VVEIADWTK |
| LDDDTIQAGVTDYSFDLTR | |

[1] Medina, M. S., Bretzing, K. O., Aviles, R. A., Chong, K. M., Espinoza, A., Garcia, C. N. G., Katz, B. B., Kharwa, R. N., Hernandez, A., Lee, J. L., Lee, T. M., Lo Verde, C., Strul, M. W., Wong, E. Y., and Owens, C. P. (2021) CowN sustains nitrogenase turnover in the presence of the inhibitor carbon monoxide, *J Biol Chem* *296*, 100501.

[2] Katz, F. E., Shi, X., Owens, C. P., Joseph, S., and Tezcan, F. A. (2017) Determination of nucleoside triphosphatase activities from measurement of true inorganic phosphate in the presence of labile phosphate compounds, *Anal Biochem* *520*, 62-67.

[3] Dilworth, M. J., Eldridge, M. E., and Eady, R. R. (1992) CORRECTION FOR CREATINE INTERFERENCE WITH THE DIRECT INDOPHENOL MEASUREMENT OF NH3 IN STEADY-STATE NITROGENASE ASSAYS, *Anal. Biochem.* *207*, 6-10.

[4] Baek, M., DiMaio, F., Anishchenko, I., Dauparas, J., Ovchinnikov, S., Lee, G. R., Wang, J., Cong, Q., Kinch, L. N., Schaeffer, R. D., Millan, C., Park, H., Adams, C., Glassman, C. R., DeGiovanni, A., Pereira, J. H., Rodrigues, A. V., van Dijk, A. A., Ebrecht, A. C., Opperman, D. J., Sagmeister, T., Buhlheller, C., Pavkov-Keller, T., Rathinaswamy, M. K., Dalwadi, U., Yip, C. K., Burke, J. E., Garcia, K. C., Grishin, N. V., Adams, P. D., Read, R. J., and Baker, D. (2021) Accurate prediction of protein structures and interactions using a three-track neural network, *Science* *373*, 871-876.
